# Exploring the effect of dynamic deformation on the tight junctions of endothelial cells

**DOI:** 10.1101/2025.09.23.677978

**Authors:** D Jahagirdar, R Kumar, A Majumder

**Affiliations:** Department of Chemical Engineering, Indian Institute of Technology Bombay, Mumbai, India

**Keywords:** Atherosclerosis, tight junction, hyperlipidaemia, dynamic narrowing, microphysiological systems

## Abstract

Atherosclerosis (Ath) is a leading cause of cardiovascular deaths, accounting for approximately 17.1 million fatalities annually. This condition arises from plaque-induced arterial narrowing, which obstructs blood flow. Traditional animal models face ethical, economic, and translational limitations, while current in vitro models often oversimplify plaque formation by using fixed channels and neglecting the dynamic effects of narrowing on endothelial junctions. The study addresses the limitations by developing a microfluidic device that mimics human-relevant stenosis with dynamic channel narrowing under continuous perfusion. Endothelial cells, which are crucial for maintaining barrier integrity through tight junctions, are subjected to physiological shear stress and hyperlipidaemic conditions by administering free fatty acids. The model evaluates cell alignment, junctional stability, and disease progression under mechanically deformed static conditions. To validate the translational potential of this system, ROS scavenging drug, quercetin is tested for its ability to reverse oxidative stress and barrier dysfunction. This innovative microphysiological system offers a robust platform for investigating endothelial dysfunction in Ath and provides a valuable alternative to animal models for cardiovascular drug testing. By integrating both biochemical and mechanical cues in a controllable environment, this model aims to enhance our understanding of Ath and facilitate the development of more effective therapeutic strategies.

## Introduction

Cardiovascular disease remains a global health burden, with atherosclerosis as its leading cause. In 2021, over half a billion people were affected, and more than 20 million deaths were attributed to cardiovascular conditions (1–3). Traditionally, animal models ranging from rodents to non-human primates have served as the gold standard for studying atherosclerosis and testing therapies (5–8). However, these models suffer from significant drawbacks including interspecies differences, long induction times, high costs, and limited throughput, making them inefficient for translational research (9–12).

Atherosclerosis is an inflammatory disease initiated by lipid accumulation within the endothelium, progressing through endothelial dysfunction, smooth muscle cell activation, and plaque formation (13). The mechanical microenvironment plays a key role in this process: plaque geometry, stiffness, and vessel narrowing evolve dynamically over time, while wall shear stress (WSS) from blood flow critically influences endothelial permeability and plaque stability (14). Rupture of unstable plaques leads to atherothrombosis, often resulting in fatal outcomes. Despite this knowledge, existing *in vitro* models primarily rely on predefined geometries or fixed deformations. Such systems capture only static aspects of stenosis and do not account for the real-time strain buildup and disturbed flow that occur during plaque progression. As a result, the dynamic interplay between mechanical strain, shear stress, and endothelial dysfunction remains poorly understood (15–17).

Endothelial cells are highly responsive to fluid shear stress and cyclic strain, which regulate barrier function, ROS production, and inflammatory signalling. To truly mimic atherosclerosis progression, there is a critical need for dynamic, deformable *in vitro* platforms that couple biochemical triggers with evolving mechanical cues (16,17). A microfluidic atherosclerosis-on-chip with tunable stenosis would enable real-time monitoring of endothelial responses under physiologically relevant strain and flow conditions, bridging the gap between static models and *in vivo* pathophysiology while offering a high-throughput, human-relevant tool for mechanistic studies and drug validation.

Lipid molecules, particularly oleic and palmitic acid, play a crucial role in the body’s metabolic processes (18). However, when present in elevated levels in the bloodstream, they can pose significant risks to the endothelial cell layer lining the arteries. Oleic acid, a monounsaturated fatty acid, can deposit within endothelial cells, leading to increased rigidity and compromised cellular function (19). This deposition disrupts the integrity of the endothelial monolayer, a critical barrier that maintains vascular homeostasis. The rigidity induced by oleic acid accumulation impairs the endothelial cells’ ability to respond to physiological stimuli, thereby promoting the formation of atherosclerotic plaques beneath the endothelial layer (20,21). These plaques, composed of lipids, cholesterol, and cellular debris, further exacerbate endothelial dysfunction and contribute to the narrowing and hardening of the arteries, a condition known as atherosclerosis. The interplay between elevated oleic acid levels and endothelial cell rigidity underscores the importance of maintaining lipid homeostasis to prevent vascular complications. Understanding the mechanisms by which oleic acid influences endothelial cell behavior is essential for developing therapeutic strategies aimed at mitigating the adverse effects of lipid accumulation and preserving cardiovascular health.

The rigidity induced by lipid accumulation, such as oleic acid, within endothelial cells significantly impacts their barrier function by disrupting tight junctions. Tight junctions are crucial for maintaining the selective permeability of the endothelial monolayer, and zona occludens 1 (ZO-1) is a key protein in this complex. ZO-1 interacts with other tight junction proteins to form a continuous seal that regulates the passage of molecules between endothelial cells (22). When endothelial cells become rigid due to lipid deposition, the structural integrity of tight junctions is compromised. This rigidity alters the spatial organization and expression of ZO-1, leading to weakened cell-cell adhesion and increased paracellular permeability. As a result, the endothelial barrier becomes more permeable, allowing harmful substances to infiltrate the arterial wall and promoting inflammation and atherosclerosis. The disruption of ZO-1 and other tight junction proteins not only impairs the barrier function but also affects the signaling pathways involved in maintaining endothelial homeostasis (23). Understanding the mechanisms by which cellular rigidity disrupts tight junctions and ZO-1 function is essential for developing therapeutic strategies to preserve endothelial integrity and prevent vascular diseases.

Static in vitro models and animal models have significant limitations when it comes to accurately replicating human physiological conditions. Static in vitro models often fail to mimic the dynamic environment of living tissues, lacking the mechanical forces and biochemical gradients present in vivo. Similarly, animal models, while providing a more complex biological context, do not always accurately represent human biology due to species-specific differences (24,25). These shortcomings highlight the need for systems that can integrate both biochemical and mechanical cues in a controllable manner. Such systems can better simulate the dynamic interactions within human tissues, providing more relevant and reliable data for biomedical research and therapeutic development.

## Materials and methods

### Cell culture

Primary human umbilical vascular endothelial cells (HUVEC) were procured from Himedia and cultured in the HiEndoXL medium. The cells were maintained at 37°C with 5% CO2 and 90% humidity. The medium replenishment was performed after every 48h until 70% confluency. Cells were utilized until passage number 7.

### Reactive oxygen species assay

To measure reactive oxygen species (ROS) levels, endothelial cells are incubated with 21 µM DCFDA (2’,7’-dichlorofluorescin diacetate) in a serum-free medium for 30 minutes at 37°C. After incubation, the cells are washed with PBS to remove excess DCFDA. The fluorescence intensity, which correlates with ROS levels, is then measured using a fluorescence microscope under FITC channel.

### Cell viability assay

To assess cell viability, endothelial cells are incubated with 2 µM Calcein AM and 4 µM Propidium Iodide (PI) in PBS for 30 minutes at 37°C. Calcein AM stains live cells green, while PI stains dead cells red. After incubation, the cells are washed with PBS to remove excess dyes. The fluorescence is then observed using a fluorescence microscope, with live cells emitting green fluorescence and dead cells emitting red fluorescence.

### Oil red O assay

To assess lipid accumulation, endothelial cells are fixed with 4% paraformaldehyde for 15 minutes at room temperature. After fixation, the cells are washed with PBS and stained with Oil Red O solution for 30 minutes. The cells are then washed with 60% isopropanol to remove excess stain. Lipid droplets within the cells will appear red under a microscope, indicating lipid accumulation.

### Immunofluorescence

The cells after the experiment were gently washed with PBS and fixed with 4% PFA at 4°C for 15 mins followed by PBS rinsing. The cells were further permeabilized using 0.1% tritonX100 for 15 mins at RT. The culture was blocked with 2% BSA for 1h at RT followed by an overnight incubation with ZO-1 antibody (5ug/ml) in 0.1% BSA. The samples were then treated with secondary antibody (GAM) Alexa Fluor 488 (1:500) for 1.5h at RT and further stained with HOECHST 33342 (1:3000) for 15 mins. The cells were observed under confocal microscope.

## Results

### Device design, fabrication and computational simulation

The aim of the study was to mimic the progressive stenosis observed in human arteries, the authors developed a PDMS-based static device with a built-in deformable surface. The fabrication involved creating a two-layer structure consisting of a PDMS block and a thin elastic layer engineered to retain air and generate a controllable pressurized cavity. This cavity was strategically aligned with a boundary to retain the culls and culture media above, such that inflation of the cavity caused local deformation of the PDMS sheet that closely reflects the in vivo plaque area as shown in **Fig 1A**. The design not only enabled dynamic narrowing of the surface but was also optimized for real-time microscopic visualization following endothelial cell seeding. This architecture provided a versatile platform to mimic plaque-induced stenosis and to directly observe endothelial responses under varying degrees of deformation. The side view schematics of the device design with cells prior to and after deformation are shown in **Fig 1B**, whereas the top view of the fabricated device is shown in **Fig 1C**.

**Fig 1:**
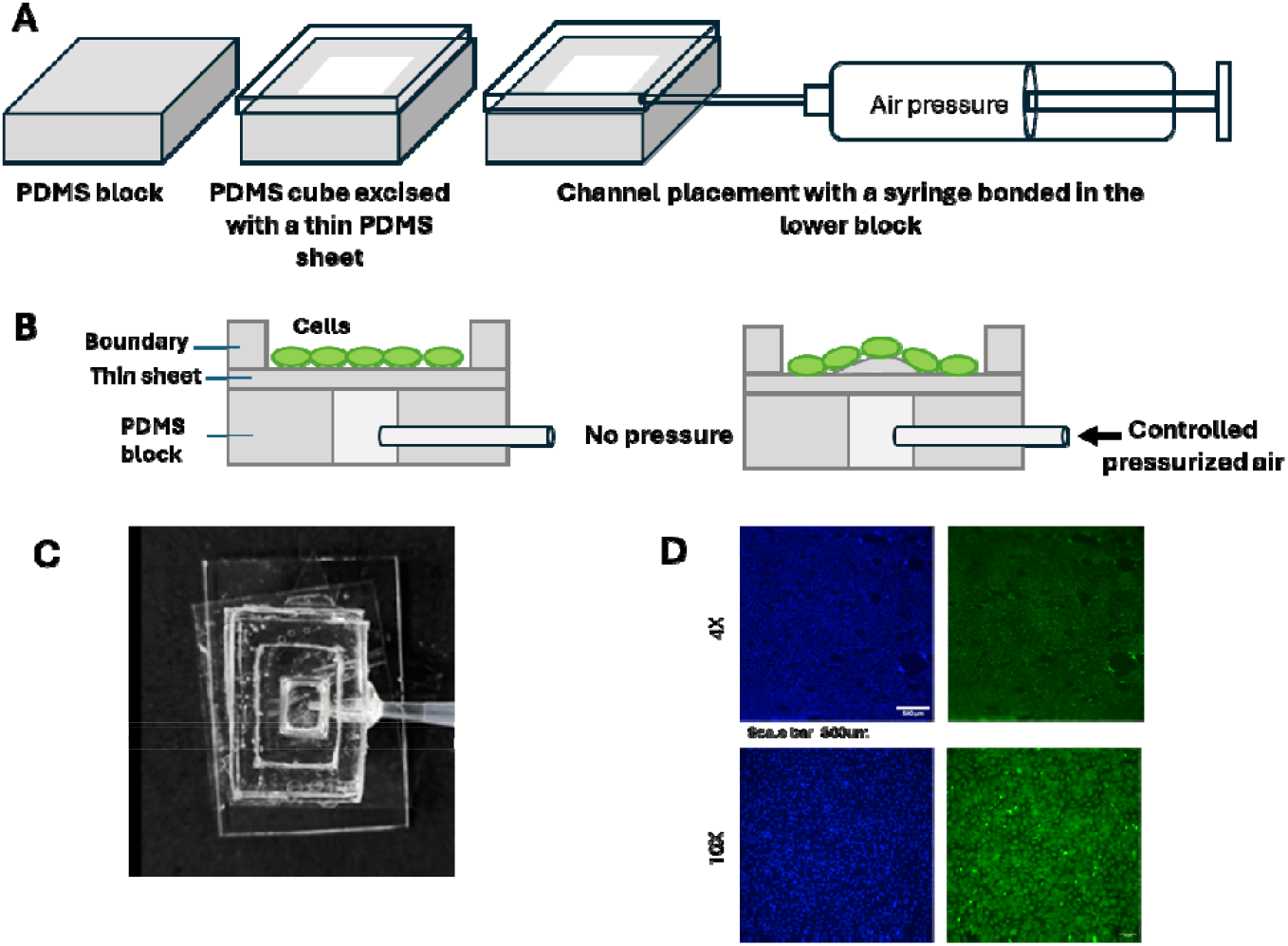
(A & B) Schematic representation of device fabrication and deformation. (C) Top view of the device. (D) Endothelial monolayer stained with Hoechst (blue) and calcein (green) at 4X and 10X.

To study the effect of the endothelial tight junction the device design successfully caters to the fundamental biology of the endothelial barrier properties. The design rationale was to create a device that could accurately replicate the mechanical environment of stenotic arteries, allowing for the study of endothelial cell behaviour under conditions that closely mimic those found *in vivo*. By using a PDMS-based structure, the authors were able to create a robust device, capable of withstanding repeated cycles of deformation without losing its structural integrity. The inclusion of a pressurized cavity allowed for precise control over the degree of deformation, enabling the researchers to study the effects of varying levels of stenosis on endothelial cell function. Endothelial cell seeding density was standardized, ensuring that cells adhered uniformly and progressed to form a confluent monolayer with 10e5 cells/cm2. The integrity and viability of this monolayer were verified using live/dead staining assays, which confirmed that optimized densities supported consistent coverage and minimal cell death. To ensure complete endothelialisation of the device surface, viability assay was performed using Hoechst (blue) and calcein (green) as depicted in **Fig1D**.

### Optimization of Disease Induction

To establish a robust *in vitro* disease platform that mimics the early stages of atherosclerosis, endothelial dysfunction induction was standardized using oleic acid (OA) treatment in a 2D culture system of human endothelial cells. The rationale was to recreate the lipid-induced oxidative stress commonly observed in vascular endothelium exposed to hyperlipidaemic conditions. For optimization, a series of dose- and time-dependent experiments wherein endothelial cells were exposed to varying concentrations of OA and monitored at 24, 48, and 72 h. Intracellular lipid deposition was quantified using the Oil Red O assay (**Fig 2A**), while oxidative stress levels were measured via DCFDA fluorescence assay as shown in **Fig 2B**. The results demonstrated a clear concentration- and time-dependent increase in both lipid accumulation and reactive oxygen species (ROS) generation. Importantly, an optimized OA concentration that required a relatively lower dose and shorter exposure time yet yielded maximal physiological effect was chosen for inducing disease-like characteristics. This optimization step was critical to minimize cytotoxicity while reliably inducing disease-like phenotypes.

**Fig 2:**
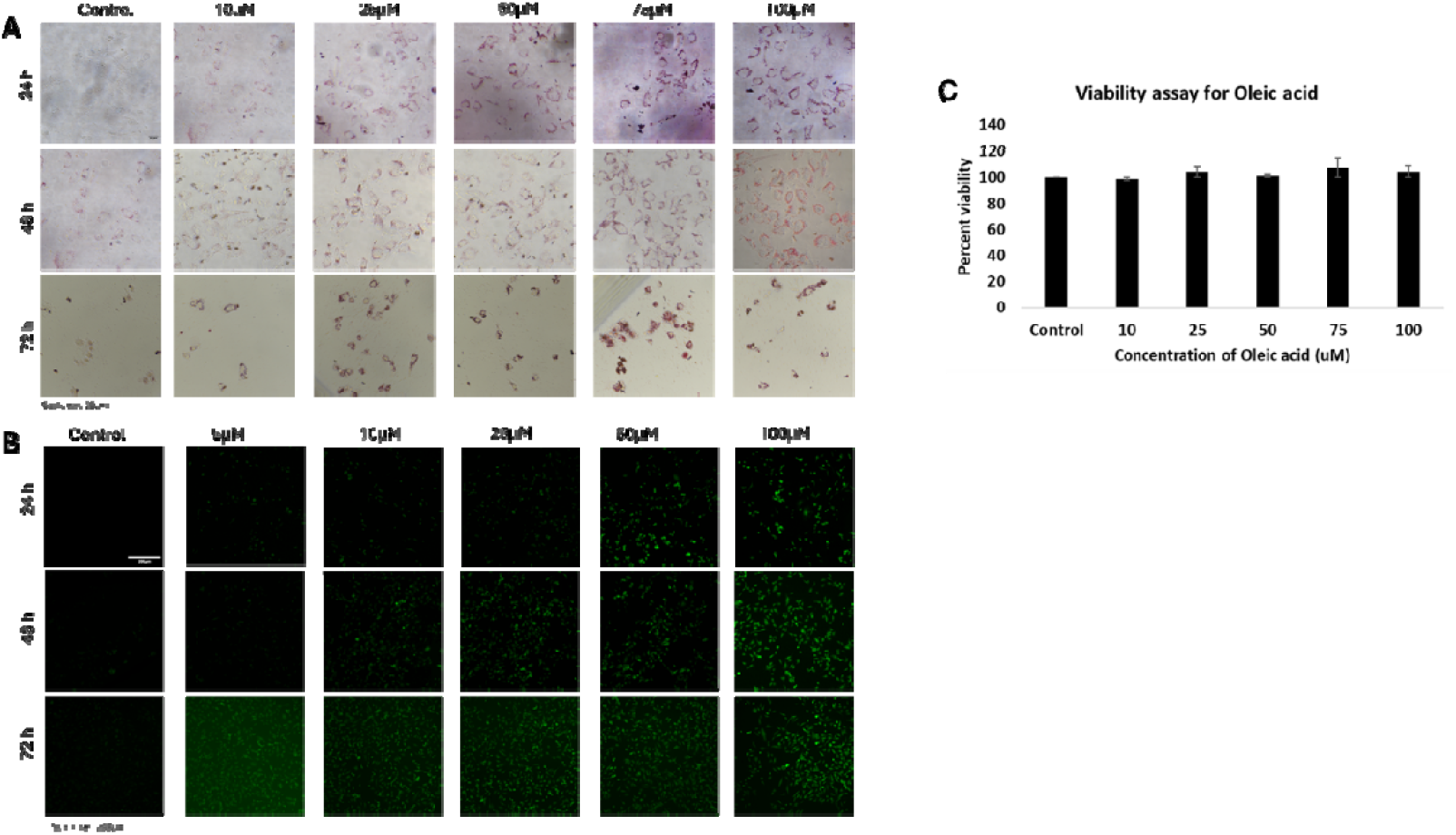
Disease induction (A) Intracellular lipid accumulation using oil red O and (B) Reactive oxygen species screened via DCFDA assay with time (24-48 h) and dose (5-100 µM) dependent study of oleic acid (OA). (C) MTT study of HUVEC with varied concentration of OA.

The aim was to induce stress without causing cell death, therefore the concentration range selected was significantly lower than the IC50 of OA. Viability assay was conducted to screen the effect of different concentration on the HUVEC for a period of 24h. The results (**Fig 2C**) demonstrate an average viability of 98% along all the OA treated conditions. The concentration of 25 µM at 48 h was selected to induce lipidemic stress in the endothelial cells for further experiments. Time and concentration were selected on the basis of minimal toxicity under maximal efficacy.

### Reversal of the disease by quercetin

Once the OA-based disease model was established, a therapeutic 2D validation using quercetin, a natural antioxidant known for its ROS-scavenging properties was performed. Prior to co-treatment experiments, the quercetin concentration was optimized via DCFDA assay to assess its direct impact on endothelial cells in the absence of OA (**Fig 3A**). With the increase in the concentration of the quercetin, the fluorescent intensity is observed to decrease till 50 µM for the duration of 24 h. The cells in 100 µM were observed to circularize and therefor the concentration was not considered for further experiments.

**Fig 3:**
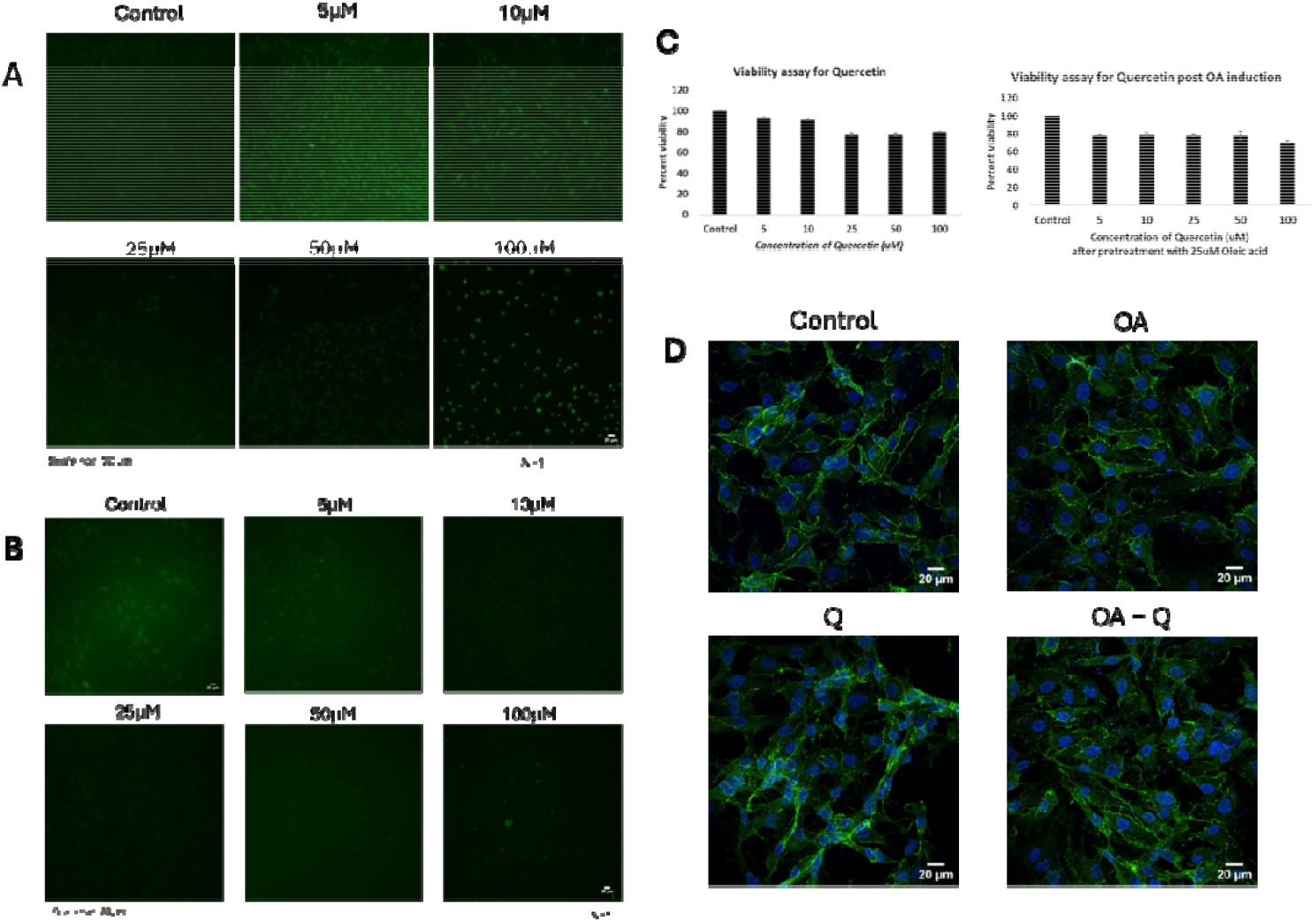
ROS expression by DCFDA (green) upon administration of (A) quercetin for 24 h and (B) Endothelial cells pretreated with 25 µM OA followed by quercetin for 24 h. (C) Viability graph via MTT for quercetin and OA pretreated cells and quercetin. (D) ZO-1 expression of the untreated cells (control), only OA (25 µM for 48 h), only quercetin (Q 25 µM for 24 h), and OA followed by quercetin (OA + Q) (N=1).

Reactive oxygen species were also assessed for the cells exposed to 25 µM of OA for 48 h. From **Fig 3B**, it is observed that quercetin was able to best quench the oxidative stress caused by 25 µM OA at the concentration of 25 µM. Through MTT assays individually conducted for quercetin and OA-induced (25 µM) quercetin (**Fig 3C**), it signifies the non-toxic effect of both the biochemical entities, and a working range that avoided cytotoxicity while maintaining protective antioxidant effects was determined. Subsequently, endothelial cells were pretreated with OA for 48h to induce lipid accumulation and oxidative stress, followed by 24h of quercetin administration. This sequential design enabled direct evaluation of quercetin’s efficacy in reversing OA-induced dysfunction. The outcomes indicated that quercetin effectively reduced intracellular ROS levels and partially restored cell viability under stress conditions, confirming its role in alleviating oxidative damage. To understand the effect of these treatment conditions on the barrier properties, the tight junction protein, ZO-1 was screened to understand the expression. **Fig 3D** depicts the difference in the continual expression of the tight junction protein expressed by the HUVEC. The immunofluorescence results depicts change in the expression in the case of OA and OA treated with quercetin in comparison with the control and only quercetin treated cells.

### Establishing deformation-on-chip

In parallel with biochemical disease induction, dynamic deformable device was used to recapitulate the progressive stenosis encountered *in vivo* and to evaluate its impact on endothelial barrier integrity. A device configuration shown in **Fig 1A** was fabricated to establish proof-of-concept and systematically optimize experimental conditions. Once the monolayer was established, controlled deformation was introduced to mimic stenotic strain and subsequently examined the organization of tight junction proteins (ZO-1). The results (**Fig 4**) indicated that deformation disrupted junctional continuity, validating that mechanical strain alone is sufficient to impair barrier function, even in the absence of shear stress.

**Fig 4:**
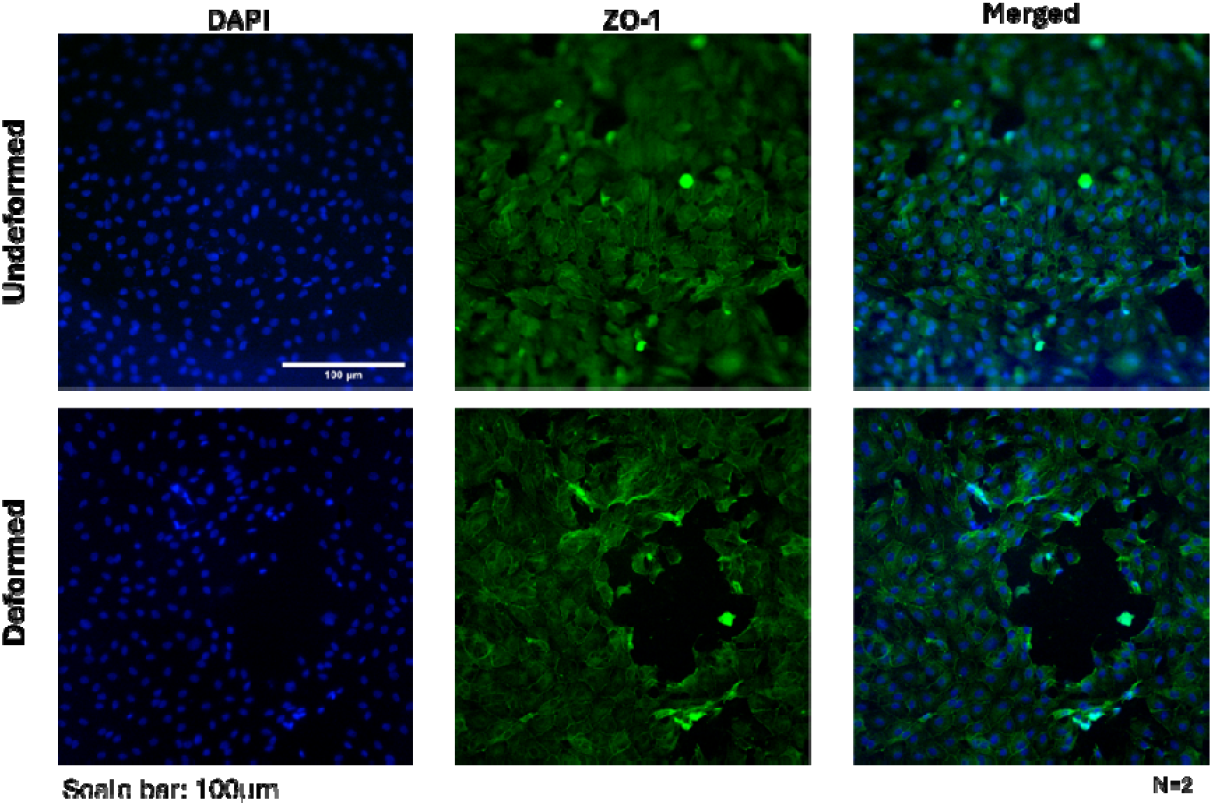
ZO-1 expression of HUVEC monolayer on the device in undeformed and deformed condition, with nucleus (blue), ZO-1 (green) (N=2).

The device provides valuable insights into the mechanisms underlying plaque formation and progression and could potentially lead to the development of new therapeutic strategies for the treatment of cardiovascular diseases. Together, the fabrication and simulation pipelines provide a robust methodology to bridge experimental vascular models with physiological pathomechanics. Importantly, this integrated approach not only validates the functional design of the deformable device but also lays the foundation for quantitatively probing the synergistic impact of shear stress and mechanical strain on endothelial dysfunction, thereby offering a more faithful representation of atherosclerotic pathophysiology in a controlled setting.

## Discussion

The deformable microfluidic device provides a physiologically relevant platform to investigate the role of mechanical stress in endothelial dysfunction. Mechanical strain on the arterial wall is a critical factor in disrupting endothelial tight junctions, a key early event in atherosclerosis that compromises barrier integrity and facilitates lipid infiltration and inflammatory cell recruitment. By allowing dynamic and tunable deformation, the PDMS-based system enables systematic study of how different magnitudes of stress contribute to junctional instability and plaque initiation.

To advance physiological relevance, a cylindrical device with continuous perfusion is currently under study. This configuration replicates both shear forces and dynamic narrowing observed in human arteries, offering a closer mimicry of in vivo vascular conditions. Unlike flat or static systems, this integrated platform will capture the combined effects of mechanical deformation, flow-induced shear, and hyperlipidaemic conditions on endothelial function. Importantly, such a design holds significant promise as a more representative system for high-throughput drug screening, where candidate formulations can be systematically tested for their ability to prevent or reverse endothelial dysfunction.

Parallel optimization using the OA-induced endothelial dysfunction model has validated a reproducible disease phenotype characterized by lipid accumulation and oxidative stress. The system has also demonstrated responsiveness to pharmacological agents such as quercetin, reinforcing its translational utility. Together, these approaches establish a solid foundation for exploring the interplay of biomechanical and biochemical cues in atherosclerosis and pave the way toward scalable platforms for mechanistic studies and therapeutic testing.

## Conclusion

In conclusion, the development of deformable microfluidic devices represents a significant step forward in modeling the biomechanical and biochemical drivers of atherosclerosis. By enabling controlled and dynamic wall deformation, these systems capture the critical role of mechanical stress in disrupting endothelial tight junctions, a process central to barrier dysfunction, lipid infiltration, and plaque progression. The complementary OA-induced disease model further establishes a reproducible and pharmacologically responsive platform that recapitulates key features of hyperlipidemia, including lipid accumulation and oxidative stress. Together, these advances provide a strong foundation for dissecting the interplay between mechanical strain, and metabolic stress in the pathogenesis of vascular disease. Importantly, the transition toward a cylindrical perfused device will enhance physiological relevance by integrating shear stress with progressive luminal narrowing, more closely mimicking in vivo arterial conditions. This evolution not only strengthens the biological insights that can be derived but also positions the system as a scalable and versatile tool for high-throughput drug screening. Future integration of disease-induced endothelial cells will further refine the model and extend its translational impact. Overall, this work lays the groundwork for a robust, human-relevant platform to study atherosclerosis and evaluate therapeutic interventions beyond the limitations of animal models.

